# Mouse Embryonic Stem Cell Pluripotency Factors Regulate RNA Methylation

**DOI:** 10.1101/2023.02.23.529801

**Authors:** Laura J. Sedivy, Gabrielle Brandt, Aspen L. Martin, Hannah M. Abroe, Christopher J. Phiel

**Affiliations:** Department of Integrative Biology, University of Colorado Denver, Denver, CO 80204

## Abstract

The pluripotency of embryonic stem cells (ESCs) is actively promoted by a diverse set of factors, including leukemia inhibitory factor (LIF), glycogen synthase kinase-3 (Gsk-3) and mitogen-activated protein kinase kinase (MEK) inhibitors, ascorbic acid, and α-ketoglutarate. Strikingly, several of these factors intersect with the post-transcriptional methylation of RNA (m^6^A), which has also been shown to play a role in ESC pluripotency. Therefore, we explored the possibility that these factors converge on this biochemical pathway to promote the retention of ESC pluripotency. Mouse ESCs were treated with various combinations of small molecules, and the relative levels of m^6^A RNA were measured, as well as the expression of genes marking naïve and primed ESCs. The most surprising result was the discovery that replacing glucose with high levels of fructose pushed ESCs to a more naïve state and reduced m^6^A RNA abundance. Our results suggest a correlation between molecules previously shown to promote the retention of ESC pluripotency and m^6^A RNA levels, strengthening a molecular connection between reduced m^6^A RNA and the pluripotent state, and provides a foundation for future mechanistic studies on the role of m^6^A and ESC pluripotency.

## Introduction

There are over 100 different chemical modifications that are found on RNA molecules, and the biological role of many of these modifications is unknown. One modification discovered over 40 years ago, methylation of adenosine bases at the C6 position (referred to as m^6^A), is the most common internal (non-cap) mRNA modification (Wei *et al*., 1975, 1976; Wei and Moss, 1977). The functional importance of m^6^A bases in mRNA proved difficult to study, and only recently has the significance of m^6^A begun to emerge (Pan, 2013; Fu *et al*., 2014; Meyer and Jaffrey, 2014; Ries *et al*., 2019). Enrichment of m^6^A followed by next-generation sequencing (m^6^A-seq) has revealed a great deal of information about the precise mRNAs that contain m^6^A, as well as their abundance and distribution along mRNAs (Dominissini *et al*., 2012; Meyer *et al*., 2012). Importantly, enzymes have been identified that add the m^6^A tags (Mettl3) (Liu *et al*., 2014) and remove the m^6^A tags (FTO and Alkbh5) (Jia *et al*., 2011; Zheng *et al*., 2013), as well as proteins whose function is to recognize and bind to the m^6^A tags (YTHDF, IGF2BP) (Wang *et al*., 2014a; Wang *et al*., 2015). The discovery of proteins that control m^6^A mRNA levels has allowed for studies to determine the biological consequences of enhancing or depleting m^6^A tags on mRNA. It has recently been reported that YTHDF proteins and IGF2BP proteins have essentially opposite functions with respect to the effects on mRNA modified by m^6^A. For example, YTHDF2 binding to m^6^A promotes mRNA decay (Wang *et al*., 2014a), while IGF2BP binding stabilizes m^6^A modified mRNAs (Huang *et al*., 2018). Therefore, an important conclusion is that it is currently not possible to infer the consequence of m^6^A modifications of mRNA by simply detecting the presence of methylated adenosines.

RNA methylation has been shown to play an important functional role in controlling the pluripotency of embryonic stem cells (ESCs). Specifically, reducing m^6^A RNA levels in ESCs causes the cells to remain pluripotent, even under conditions where differentiation should occur normally (Batista *et al*., 2014; Wang *et al*., 2014b; Geula *et al*., 2015). Initial characterization of this effect revealed that the mRNA for many regulators of stem cell pluripotency are normally methylated, and in the absence of Mettl3, m^6^A mRNA levels are reduced, causing the half-life of these mRNAs to be increased, presumably resulting in increased levels of protein. However, m^6^A modifications to mRNA have also been shown to affect RNA splicing, export and translation (Patil *et al*., 2018). Depending on which reader protein binds to m^6^A, opposite outcomes can result. Therefore, while reduced m^6^A mRNA levels are an important factor in controlling stem cell pluripotency, the exact reason for this effect is still unclear.

Small molecule inhibitors of glycogen synthase kinase-3 (Gsk-3) have also been shown to promote pluripotency (Sato *et al*., 2004; Bone *et al*., 2009), particularly when combined with MEK inhibitors; this has been termed the 2 inhibitor (2i) cocktail, and has been shown to promote the ground state of pluripotency for ESCs, i.e., their most naïve state (Ying *et al*., 2008). The *Gsk-3* DKO ESCs show many of the hallmarks of being naïve, with the additional facet of being resistant to differentiation, even in the absence of LIF (Doble *et al*., 2007; Faulds *et al*., 2018). Gsk-3 inhibitors have also been used to facilitate the derivation of ESCs from different strains of mice (Umehara *et al*., 2007; Ye *et al*., 2012) and different species (Buehr *et al*., 2008; Li *et al*., 2008). Furthermore, genetic deletion of *Gsk-3α* and *Gsk-3β* in mouse ESCs results in cells that are unable to differentiate (Doble *et al*., 2007). Taken together, these data point to a fundamental role for Gsk-3 inhibition in promoting ESC pluripotency. Recently, we found a direct biochemical connection between Gsk-3 and m^6^A RNA modifications. The RNA demethylase FTO is phosphorylated by Gsk-3, targeting FTO for ubiquitination and degradation by the 26S proteasome (Zhu *et al*., 2018), and that this process is impaired in *Gsk-3* DKO ESCs (Faulds *et al*., 2018). As a consequence, m^6^A RNA levels are reduced by 50% in *Gsk-3* DKO ESCs, which provides a potential mechanism for how Gsk-3 inhibitors are promoting ESC pluripotency.

The experiments described here were performed to better understand the regulation of ESC pluripotency, and to determine if factors promoting ESC pluripotency correlated with changes in m^6^A RNA abundance. We included experiments using *Gsk-3* DKO ESCs since these cells are more pluripotent than WT ESCs, and could shed some light on why the *Gsk-3* DKO ESCs are resistant to differentiation.

## Results and Discussion

### Pluripotency index and m^6^A quantification

In this study, our goal was to examine the effects of specific molecules on mouse ESC pluripotency, and to assess whether any relationship exists between pluripotency and the relative abundance levels of RNA methylation. To measure m^6^A RNA levels, we performed ELISAs using a commercially available kit, and by including a standard curve, we quantified the relative amount of m^6^A RNA in each condition. In prior m^6^A analyses, we isolated total RNA from ESCs, enriched for mRNA using oligo (dT) beads, then quantified m^6^A-modified nucleotides via UPLC (Faulds *et al*., 2018). Because this current study manipulated numerous ESC culture conditions, we wanted to streamline our m^6^A quantification without sacrificing sensitivity. Therefore, we quantified m^6^A from total RNA using RNA isolated from WT and *Gsk-3* DKO ESCs that were grown in the absence of LIF (–LIF), a condition we used in our previous study (Faulds *et al*., 2018). In that study, using enriched mRNA, we found *Gsk-3* DKO ESCs had 50% less m^6^A than WT ESCs (Faulds *et al*., 2018). In comparison, using total RNA, we saw an almost identical reduction in m^6^A in *Gsk-3* DKO ESCs – 47% (Table 1). This concordance gave us the confidence to move forward quantifying m^6^A from total RNA via ELISA in the experiments described here.

**Table 1.** Quantification of pluripotency and m^6^A RNA levels. The ratio of *Nanog:Fgf5* expression from qPCR data was calculated and represented as the pluripotency index (P_i_). For all experiments using WT ESCs, P_i_ levels are relative to the expression of *Nanog* and *Fgf5* in untreated WT ESCs, which was set to 1. Similarly, for all experiments in *Gsk-3* DKO ESCs, P_i_ levels are relative to the expression of *Nanog* and *Fgf5* in untreated *Gsk-3* DKO ESCs. m^6^A% were obtained from comparison of sample values to a standard curve.

|  | <u>P<sub>i</sub></u> | <u>% m<sup>6</sup>A</u> |
| --- | --- | --- |
| WT | 1 | 0.2458 |
| WT -Lif | 0.2 | 0.3439 |
| WT FTO | 2.8 | 0.2045 |
| WT VitC | 19.1 | 0.1838 |
| WT FTO αKG Gsk-3i MEKi | 19.1 | 0.1514 |
| WT FTO VitC αKG | 7 | 0.1174 |
| DKO | 1 | 0.1338 |
| DKO FTO | 2 | 0.1095 |
| DKO VitC | 3.2 | 0.1029 |
| DKO No Glucose | 0.02 | 0.1577 |
| DKO High G | 1 | 0.3409 |
| DKO Fructose | 1.1 | 0.1541 |
| DKO High F | 2 | 0.1113 |
| DKO FTO High F | 2.3 | 0.1015 |
| DKO VitC High F | 6.5 | 0.1232 |
| DKO αKG | 0.9 | 0.2155 |
| DKO FTO αKG | 1.8 | 0.2774 |
| DKO FTO αKG MEKi | 11.6 | 0.1742 |

To measure ESC pluripotency, we performed qPCR for both *Nanog* and *Fgf5. Nanog* is a well-characterized marker of pluripotency, and is highly expressed in naïve ESCs (Chambers *et al*., 2003; Mitsui *et al*., 2003), while *Fgf5* has been shown to be a reliable marker for slightly more differentiated primed ESCs (Pelton *et al*., 2002). To help us interpret the qPCR data, we examined the ratio of *Nanog:Fgf5* expression, a value we have termed the pluripotency index (P_i_); higher Pi values indicate more naïve ESCs, while lower P_i_ values reflect more primed ESCs. The ratio of *Nanog:Fgf5* was used because relying on the expression of *Nanog* alone, for example, only provides information about ESCs in the naïve state, but does not inform about their relationship to primed ESCs.

### WT ESCs

#### LIF

Growing WT ESCs in the absence of LIF releases them from pluripotency and the ESCs initiate differentiation into primed ESCs. This was evident from the *Nanog:Fgf5* ratio, with WT ESCs grown in the presence of LIF having a higher P_i_ than WT ESCs grown without LIF (1 vs. 0.2) (Figure 1; Table 1). The relative abundance of m^6^A RNA showed the opposite trend, with WT ESCs +LIF containing less m^6^A than WT –LIF ESCs (0.2458% vs. 0.3439%) (Table 1).

**Figure 1.**
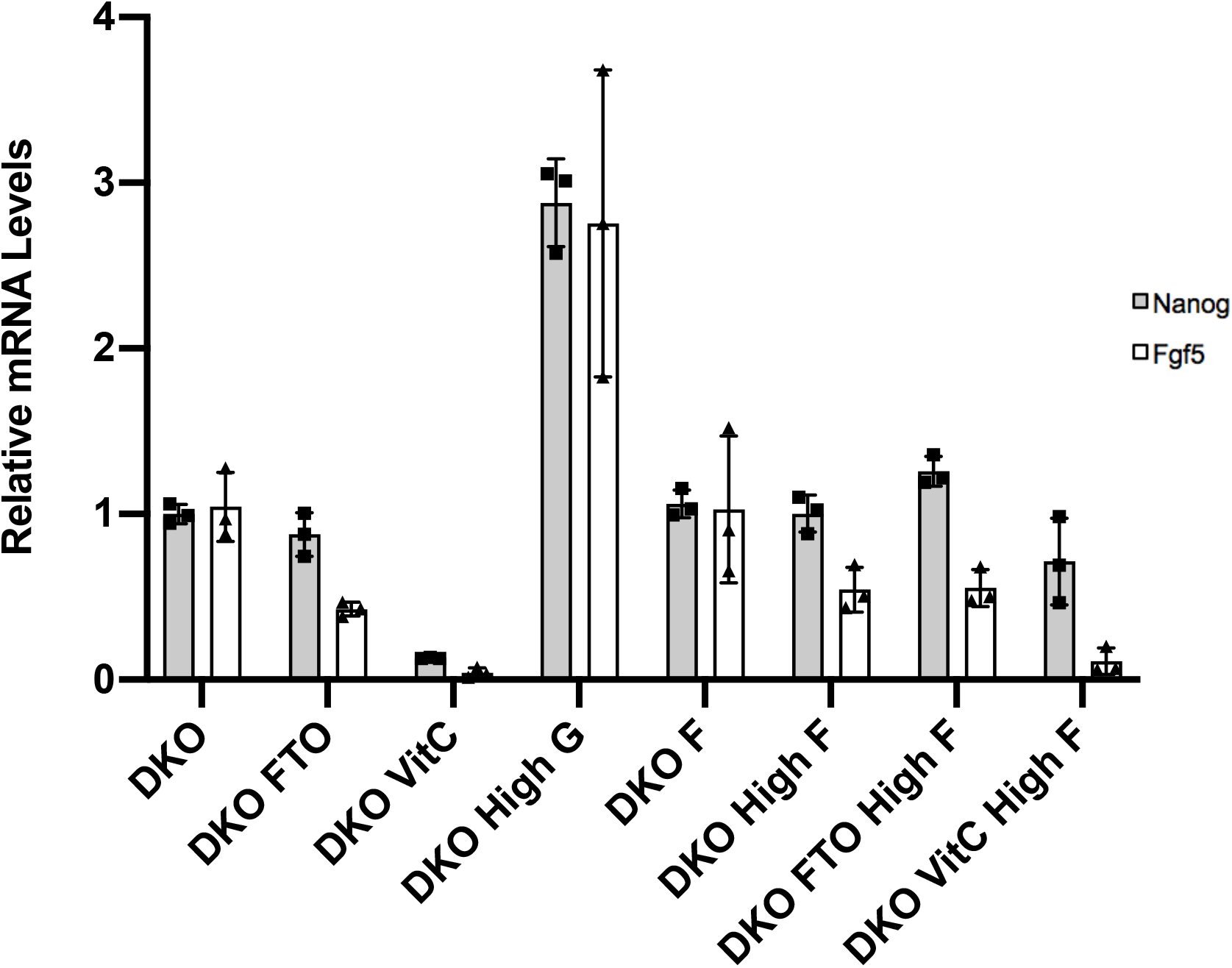
Effect of LIF, FTO, Vitamin C, α-ketoglutarate and 2i on pluripotency in WT ESCs. qPCR data showing the relative quantification (RQ) of *Nanog* and *Fgf5* gene expression in WT ESCs normalized to *Gapdh*. Each experiment was performed in triplicate, and each sample was measured in triplicate. Error bars represent standard deviation. Fold changes of *Nanog* and *Fgf5* are based on relative levels measured in WT ESCs.

#### FTO

We next assessed the effect of overexpressing the RNA demethylase FTO in WT ESCs. Transfecting ESCs with pCAGEN-FTO had the expected effect of reduced m^6^A levels (17% reduction in WT ESCs), which is identical to the effect of overexpressing FTO in HeLa cells (Jia *et al*., 2011). In addition, overexpression of FTO led to an increase in pluripotency (P_i_ of 2.8 in FTO overexpression vs. 1 in WT ESCs) (Figure 1; Table 1). Reduced m^6^A RNA levels and increased pluripotency have been noted by several groups (Batista *et al*., 2014; Geula *et al*., 2015; Bertero *et al*., 2018), and our data confirm that directly reducing m^6^A RNA levels by overexpression of FTO concomitantly increased the *Nanog:Fgf5* ratio, providing additional evidence that reduced m^6^A RNA levels is a driver of ESC pluripotency.

#### Vitamin C and α-ketoglutarate

Ascorbic acid (Vitamin C) is known to be required for FTO enzymatic activity *in vitro* (Gerken *et al*., 2007). In addition, several studies have demonstrated a role for ascorbic acid in promoting pluripotency, mostly in the context of increasing the efficiency of deriving induced pluripotent stem cells (iPSCs) (Esteban and Pei, 2012; Stadtfeld *et al*., 2012; Bar-Nur *et al*., 2014). Consistent with these studies, we found that treating WT ESCs with 50 μg/ml ascorbic acid resulted in the highest pluripotency index observed in WT ESCs (P_i_=19.1), largely due to a strong suppression of *Fgf5* expression (Figure 1; Table 1). Coincident with a high P_i_ value, we also observed a 25% reduction in m^6^A abundance (0.2458% vs. 0.1838%) (Table 1).

Another co-factor that is required for FTO activity *in vitro* is α-ketoglutarate (also referred to as 2-oxoglutarate) (Gerken *et al*., 2007). α-ketoglutarate is a co-factor for a variety of enzymes, including Tet family DNA demethylases (Tahiliani *et al*., 2009) and Jumonji-C domain family histone demethylases (Tsukada *et al*., 2006), making it a common co-factor for enzymes that demethylate RNA, DNA, and protein. It was recently reported that treating WT ESCs with a cell-permeable α-ketoglutarate, in the presence of Gsk-3 and MEK inhibitors, was sufficient to promote pluripotency (Carey *et al*., 2015). Therefore, we tested the effect of α-ketoglutarate in combination with different small molecules. Importantly, we found that in ESCs overexpressing FTO and treated with α-ketoglutarate and Gsk-3 and MEK inhibitors, pluripotency was greatly enhanced (P_i_=19.1), confirming the work of Carey and colleagues (Figure 1; Table 1). Additionally, we extended this work by demonstrating that m^6^A RNA abundance decreased 39% under these conditions (0.2458% vs. 0.1514%). This experiment suggests that the effects of α-ketoglutarate on pluripotency are at least in part through its role as a cofactor for FTO.

Characterization of FTO *in vitro* showed a requirement for *both* Vitamin C and α-ketoglutarate in demethylation assays (Gerken *et al*., 2007). Therefore, we predicted that treating WT ESCs with both Vitamin C and α-ketoglutarate should enhance m^6^A RNA demethylation and enhance pluripotency. This was indeed the case, with WT ESCs overexpressing FTO and treated with Vitamin C and α-ketoglutarate having the largest reduction in m^6^A RNA (52% reduction) and a robust increase in pluripotency (P_i_=7.0) (Figure 1; Table 1).

### Gsk-3α^-/-^; Gsk-3β^-/-^ ESCs

In addition to performing experiments on WT ESCs, we took advantage of our expertise in Gsk-3 biology to examine factors affecting pluripotency in *Gsk-3α*^-/-^; *Gsk-3β*^-/-^ double knockout (DKO) ESCs. The goal was to test if there were any cell culture conditions that resulted in *Gsk-3* DKO ESCs being more or less pluripotent, and if so, ask whether the abundance of m^6^A RNA also changed.

### FTO

We first assessed the effect of overexpressing FTO in *Gsk-3* DKO ESCs. Similar to what was observed in WT ESCs, transfecting *Gsk-3* DKO ESCs with pCAGEN-FTO reduced m^6^A RNA levels by 18% (Table 1). This was accompanied by an increase in the pluripotency index (2.0 in FTO overexpression vs. 1.0 in *Gsk-3* DKO ESCs) (Figure 2; Table 1). These data confirm the connection between reduced m^6^A RNA methylation and expression of pluripotency-related markers, and also show that these effects are similar in both WT and *Gsk-3* DKO ESCs.

**Figure 2.**
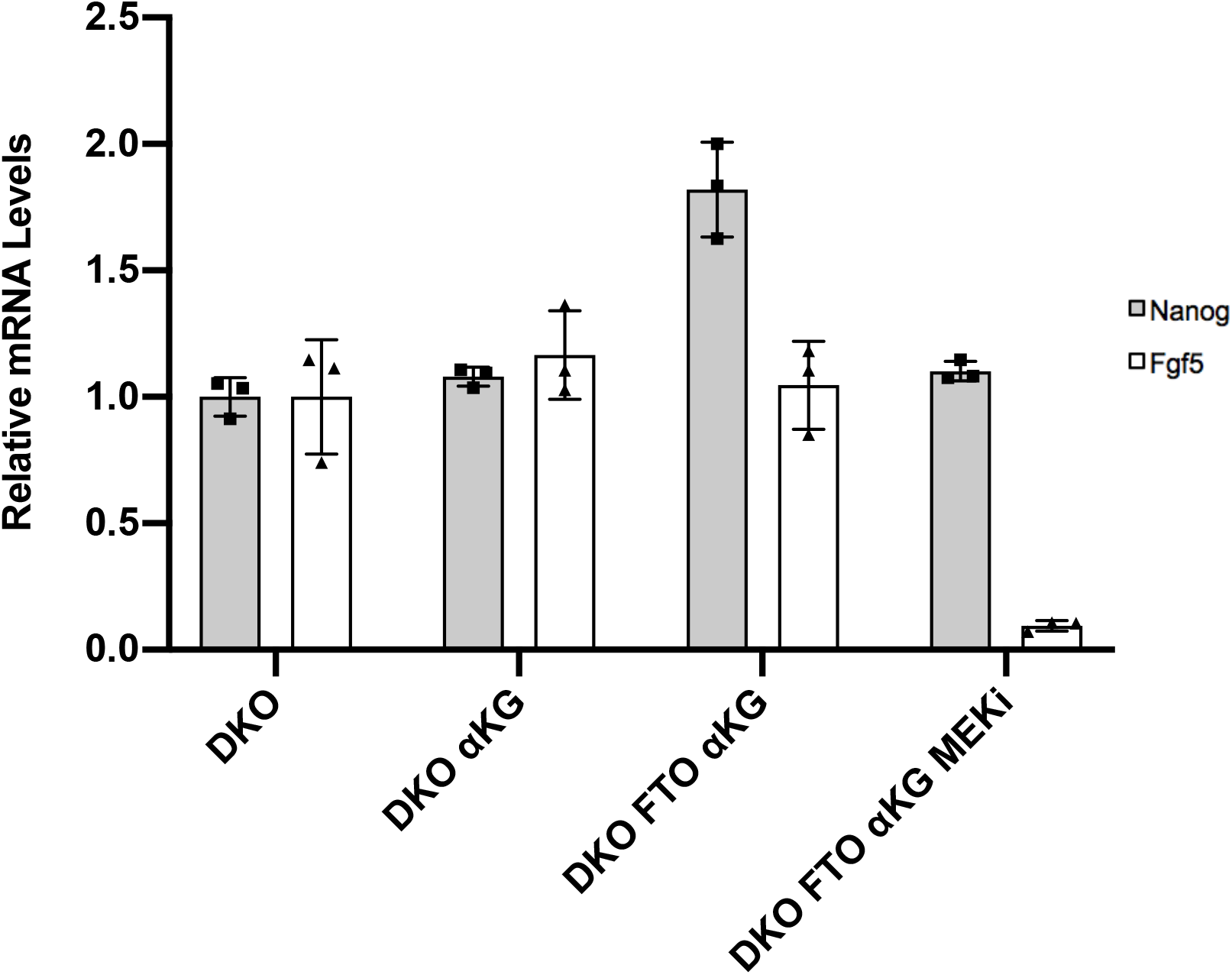
Effect of FTO, Vitamin C, and fructose on pluripotency in Gsk-3 DKO ESCs. qPCR data showing the relative quantification (RQ) of *Nanog* and *Fgf5* gene expression in *Gsk-3* DKO ESCs normalized to *Gapdh*. Each experiment was performed in triplicate, and each sample was measured in triplicate. Error bars represent standard deviation. Fold changes of *Nanog* and *Fgf5* are based on relative levels measured in *Gsk-3* DKO ESCs.

### Vitamin C and α-ketoglutarate

Hochedlinger and colleagues have previously shown that treating iPSCs with Vitamin C and a Gsk-3 inhibitor was sufficient to promote a dramatic increase in the efficiency of iPSC derivation (Bar-Nur *et al*., 2014). Therefore, we were interested to determine whether Vitamin C could further enhance pluripotency in *Gsk-3* DKO ESCs. Indeed, treatment of *Gsk-3* DKO ESCs with 50 μg/ml Vitamin C increased the pluripotency of cells already refractory to differentiation (P_i_=3.2) (Figure 2; Table 1). The increased pluripotency was accompanied by a 23% decrease in the overall abundance of m^6^A RNA (Table 1).

Based on the data obtained from WT ESCs, where α-ketoglutarate reduced m^6^A RNA levels and increased pluripotency, we expected to see similar results in *Gsk-3* DKO ESCs. Unexpectedly, α-ketoglutarate, under a variety of conditions, did not reduce m^6^A RNA abundance, but instead increased relative levels of m^6^A RNA, even when the pluripotency index was increased (Figure 3; Table 1). Taken together, these results suggest that in cells lacking Gsk-3, α-ketoglutarate promotes pluripotency, yet this is occurring without a corresponding decrease in m^6^A RNA, which was predicted based on the data from WT ESCs. Gsk-3 has many cellular functions and regulates many signal transduction pathways; perhaps the tonic loss of Gsk-3 has additional effects that make the cells unable to demethylate RNA when treated with α-ketoglutarate. Unraveling this mechanism will require further investigation.

**Figure 3.**
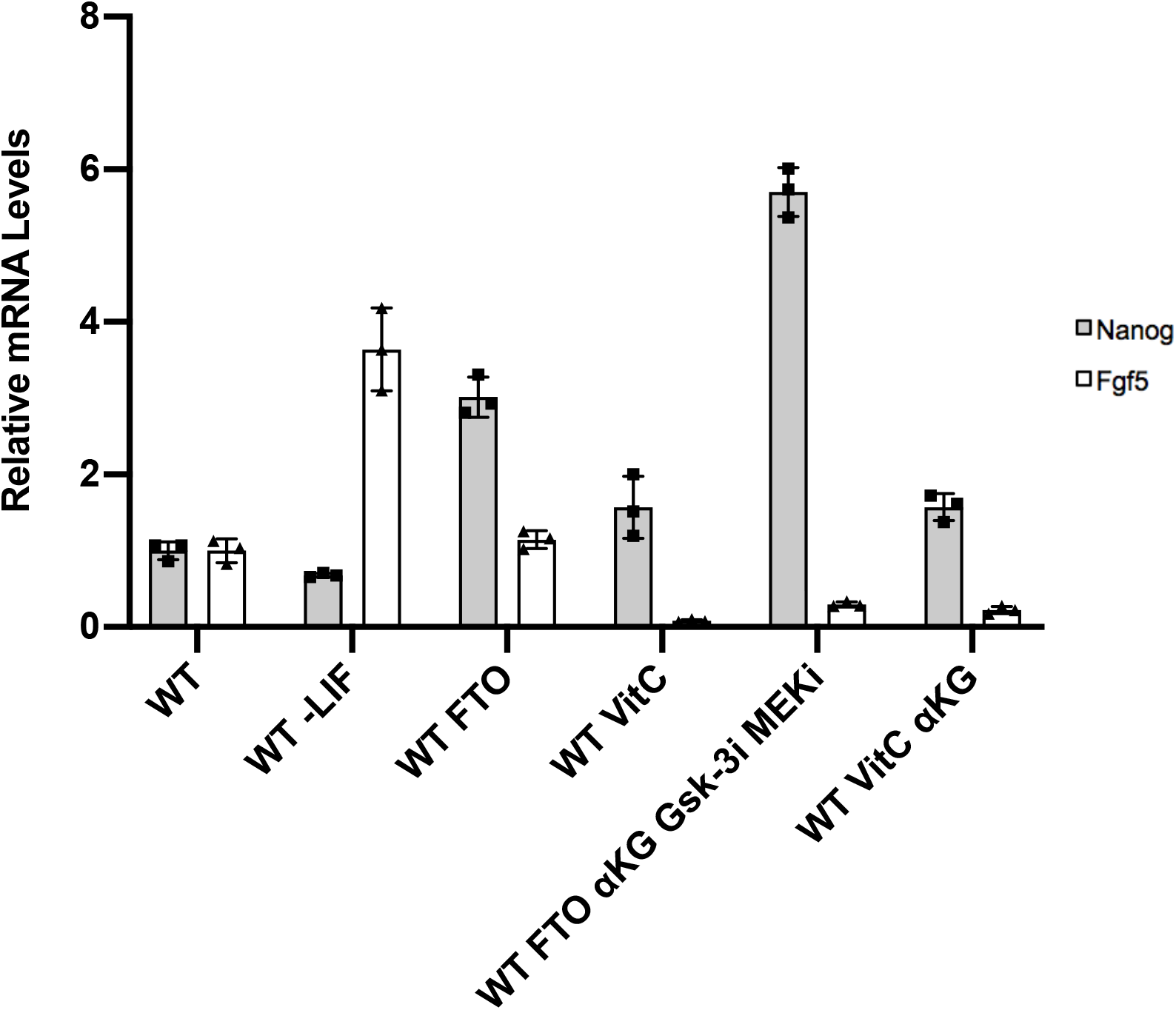
Effect of α-ketoglutarate treatment on pluripotency in Gsk-3 DKO ESCs. qPCR data showing the relative quantification (RQ) of *Nanog* and *Fgf5* gene expression in *Gsk-3* DKO ESCs normalized to *Gapdh*. Each experiment was performed in triplicate, and each sample was measured in triplicate. Error bars represent standard deviation. Fold changes of *Nanog* and *Fgf5* are based on relative levels measured in *Gsk-3* DKO ESCs.

### Cellular metabolism

Finally, we performed a set of experiments that was initiated to determine whether changes in standard ESC media could affect *Gsk-3* DKO ESC pluripotency. Our ESCs are cultured in standard DMEM containing 4.5 g/L glucose, supplemented with glutamine, pyruvate, and antibiotics. In particular, we were curious to see if altering glucose levels had an effect on pluripotency in *Gsk-3* DKO ESCs. Our initial assay was visual, plating equal numbers of cells in wells of a 24-well plate, growing Gsk-3 DKO ESCs in various media formulations, and then observing any changes in cellular morphology that might indicate an effect on pluripotency.

In one condition, we doubled the amount of glucose to 9 g/L by supplementing standard DMEM with additional glucose. This condition did not change pluripotency (P_i_=1.0) (Figure 2; Table 1), but resulted in the largest increase in m^6^A RNA abundance seen in our study (255% increase) (Table 1). Conversely, growing the ESCs in the absence of glucose actively promoted differentiation (P_i_=0.02) (Figure 2; Table 1), while also resulting in elevated m^6^A RNA levels (18% increase) (Table 1).

We then asked whether growing *Gsk-3* DKO ESCs in the presence of a different carbon source would affect the pluripotency of these cells. We selected fructose which, like glucose, is a monosaccharide; however, fructose and glucose are metabolized differently and presumably should have different effects on ESC pluripotency. We purchased glucose-free DMEM, supplemented with either 4.5 g/L fructose or 9 g/L fructose (high fructose), cultured *Gsk-3* DKO ESCs under these conditions. Surprisingly, fructose supported robust growth of *Gsk-3* DKO ESCs, with cells grown in high fructose appearing more pluripotent than ESCs grown in high glucose (Figure 4). We next analyzed the fructose effects on pluripotency markers and RNA methylation. While 4.5 g/L fructose had modest effects on m^6^A RNA (15% increase) and pluripotency (P_i_=1.1), high fructose resulted in both increased pluripotency (P_i_=2.0) and reduced m^6^A RNA levels (17% reduction) (Figure 2; Table 1).

**Figure 4.**
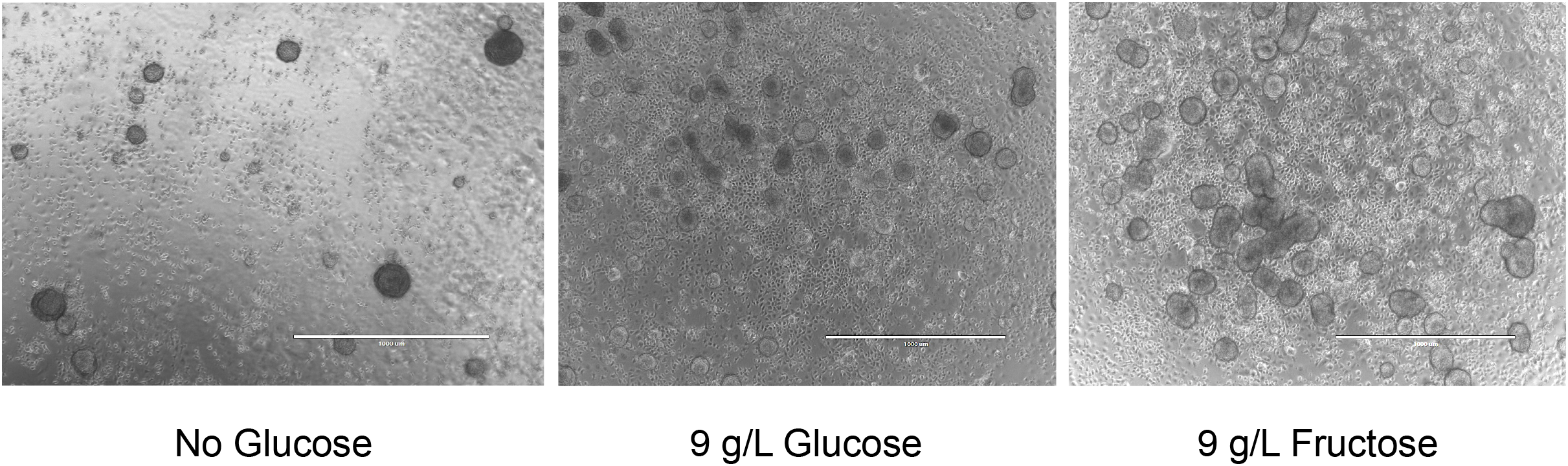
Phenotypic effects of no glucose, high glucose, and high fructose on Gsk-3 DKO ESCs. *Gsk-3* DKO ESCs were grown (A) in the absence of glucose, (B) in the presence of 9 g/L glucose, or (C) without glucose, but with 9 g/L fructose. Images were captured on an EVOS-FL microscope at 4x magnification. Scale bar represents 1000 μM.

Because we had seen increased pluripotency and reduced m^6^A RNA when *Gsk-3* DKO ESCs were overexpressing FTO or treated with Vitamin C, we asked whether those conditions synergized when cells were grown in high fructose. *Gsk-3* DKO ESCs overexpressing FTO in the presence of 9 g/L fructose resulted in m^6^A RNA abundance was the lowest that we measured (0.1015% vs. 0.1338% in standard DMEM), and the pluripotency index was 2.3 (Figure 2; Table 1). Growing *Gsk-3* DKO ESCs in Vitamin C and high fructose also resulted in increased pluripotency (P_i_=6.5) and reduced m^6^A RNA (0.1232% vs. 0.1338%) (Figure 2; Table 1).

## Conclusions

Taken together, this data provides strong evidence of a molecular connection between m^6^A RNA abundance and the pluripotency of ESCs. We found that Vitamin C promotes pluripotency in almost all conditions tested, and appears to be in part through its suppression of the transition to the primed state, as indicated by reduced expression of *Fgf5*. Furthermore, while it was expected that Vitamin C would result in reduced m^6^A RNA levels due to its role as a cofactor for FTO, to our knowledge this is the first report showing Vitamin C affects m^6^A RNA in ESCs. In addition, while the iPSC derivation study showing the potent effects of Vitamin C and a Gsk-3 inhibitor (Bar-Nur *et al*., 2014) did not examine m^6^A RNA levels, it is tempting to speculate that the effect of combined treatment with Vitamin C and Gsk-3 inhibitor was at least partially due to an effect on m^6^A RNA, with the Gsk-3 inhibitor increasing FTO protein levels and Vitamin C providing a necessary co-factor for FTO function.

The second novel observation was that replacing glucose with high levels of fructose was a strong promoter of both pluripotency and reduced m^6^A RNA abundance. Since high glucose does not have as potent an effect on pluripotency as high fructose, it is suggestive of a rate-limiting step that affects glucose metabolism but not fructose metabolism. Phosphoglucose isomerase is the enzyme that converts glucose-6-phosphate to fructose-6-phosphate in mammalian cells. Our data suggests that growing ESCs in high fructose bypasses the activity of this enzyme, allowing faster conversion to pyruvate. Based on the data gathered thus far, we can only speculate about the mechanism for the effects of high fructose on m^6^A and pluripotency, but a reasonable hypothesis is that high fructose conditions generate higher levels of α-ketoglutarate, enhancing the activity of FTO. Regardless of mechanism, it seems that replacing glucose with high fructose would be a simple and beneficial change to current ESC media formulations that could facilitate the culture of ESCs. It will be interesting to see if high fructose conditions also enhance the derivation of iPSCs.

## Materials and Methods

### Cell Culture and Transfection

Low passage, feeder-free wild-type (WT) mouse (E14K) and *Gsk-3* DKO ESCs, were grown on 0.1% gelatin-coated plates with standard DMEM containing 4.5 g/L glucose (Gibco) supplemented with 15% fetal bovine serum (HyClone), 1% non-essential amino acids (Gibco), 1% sodium pyruvate (Gibco), 1% L-glutamine (Gibco), 1% penicillin/streptomycin (Gibco) 55 μM 2-mercaptoethanol (Gibco), and 1000 units/mL recombinant leukemia inhibitory factor (LIF). Media was replenished every other day. For experiments using increased glucose (9 g/L), standard DMEM was supplemented with D-(+)-Glucose (MP Biomedicals, LLC). For experiments using a different sugar, no glucose DMEM (Gibco) was supplemented with β-D-(-)-Fructose (MP Biomedical, LLC) or D-Sucrose (Fisher). Additional experiments supplemented media with L-ascorbic acid 2-phosphate trisodium salt (Wako), dimethyl 2-oxoglutarate (Sigma Aldrich), NADH disodium salt (EMD Millipore), NADPH tetrasodium salt (EMD Millipore), PD0325901 (Selleckchem) and SB415286 (Tocris).

WT and *Gsk-3* DKO ESCs were transfected using PEI (Bartman *et al*., 2015). 1 x 10^6^ ESCs/well were resuspended in OptiMEM (Gibco) with 1800 ng of pCAGEN-FLAG-FTO, along with 200 ng of pMax-GFP. After incubation for 30 min at room temperature, cells were added to gelatin-coated 6-well plates containing complete media. Successful transfections were confirmed after 18 hours via fluorescence microscopy (Evos FL; ThermoFisher Scientific). When treating transfected cells with inhibitors, media was removed after 18 hours and replaced with fresh media containing small molecules. Cells were then harvested 24 hours later.

### RNA Isolation, cDNA Synthesis and Quantitative PCR

1 x 10^6^ ESCs/well were plated on gelatin-coated 6-well plate. RNA was isolated using TRIzol reagent (ThermoFisher Scientific) and extracted using Direct-zol RNA Miniprep columns (Zymo) following the manufacturer’s protocol. RNA was quantified using a Nanodrop One^C^ (ThermoFisher Scientific). 2 μg of total RNA was used to synthesize cDNA using the High Capacity Reverse Transcriptase kit (Applied Biosystems) according to the manufacturer’s protocol. The amount of input RNA used was kept constant for each RT reaction. Reactions were run on a StepOne Real-Time PCR System (Applied Biosystems) using PrimeTime gene expression master mix (IDT) and PrimeTime qPCR assays, *Nanog* (Mm.PT.58.23510265) and *Fgf5* (Mm.PT.58.17838080) (IDT). Three biological replicates and three technical replicates were used for WT and *Gsk-3* DKO ESCs. All threshold cycle (Ct) values were normalized to a mouse *Gapdh* endogenous control (Mm.PT.39a.1) (IDT), and relative quantification was calculated from the median Ct value.

### m^6^A Quantification

m^6^A levels were quantified using the EpiQuik m^6^A RNA Methylation Quantification Kit (Epigentek) following the manufacturer’s instructions. Briefly, 200 ng of total RNA from ESCs was bound to wells in a clear 96-well plate via RNA high binding solution, then incubated at 37°C for 90 minutes. Capture antibody was incubated at RT for 60 minutes, washed 3x, incubated with detection antibody for 30 minutes at RT, washed 3x, and finally incubated with enhancer solution for 30 minutes at RT. After washing 5x, colorimetric detection of the signal was obtained by incubating with a developer solution and then a stop solution, followed by detection at OD 450 nm using an EnSpire plate reader (PerkinElmer). A standard curve of m^6^A RNA (provided with kit) was included on each plate, in duplicate, ranging from 0.02 ng to 1 ng. The OD readings were then used to calculate the relative quantification using the following formula:

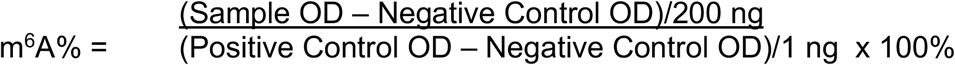

## Abbreviations

m^6^A: methyladenosine
ESCs: embryonic stem cells
Gsk-3: glycogen synthase kinase-3
DKO: double knockout
LIF: leukemia inhibitory factor
P_i_: pluripotency index
Fgf5: fibroblast growth factor 5

## Acknowledgements

We thank Xiaojun Ren for his insightful comments and suggestions. This research is supported by NIH R15GM119103 (C.J.P.)

